# cir-DNA sequencing revealed the landscape of extrachromosomal circular DNA in articular cartilage and the potential roles in osteoarthritis

**DOI:** 10.1101/2022.01.27.477985

**Authors:** Wei Xiang, Tongyi Zhang, Song Li, Yunquan Gong, Xiaoqing Luo, Jing Yuan, Yaran Wu, Xiaojing Yan, Yan Xiong, Jiqin Lian, Guangyu Zhao, Changyue Gao, Liang Kuang, Zhenhong Ni

## Abstract

**Objective:** Extrachromosomal circular DNA (eccDNA) has been shown to be involved in several physiological and pathological processes including immunity, inflammation, aging and tumor. However, the expression of eccDNA in cartilage has been not reported until now. In this study, we aimed to investigate the landscape of eccDNA in articular cartilage and analyze the potential roles in Osteoarthritis (OA).

**Methods:** The samples of articular cartilage were obtained from total knee arthroplasty (TKA) donors with OA. The mtDNAs (mitochondrial DNA) and the linear DNAs from chondrocytes of articular cartilage were removed. Then the eccDNAs were enriched for cir-DNA sequencing. After quality control evaluation, we systematically revealed the identified eccDNA data including size distribution, the size range and sequence pattern. Moreover, we explored and disscussed the potential roles of eccDNA in OA via motif analysis and GO/KEGG pathways analysis.

**Results:** The chondrocytes from OA cartilage contained an abundance of eccDNAs, which was termed as OC-eccDNAs (OA Cartilage-derived eccDNA). The characteristics of OC-eccDNAs were tissue specific, including the distribution, the size range and sequence pattern. Moreover, the functional analysis indicated that eccDNA may be involved in the homeostasis maintenance of chondrocytes and participated in the process of OA.

**Conclutions:** Our data firstly showed the landscape of eccDNA in articular cartilage, and preliminarily indicated the potential roles of eccDNA in OA.

## Introduction

Osteoarthritis (OA) is a common degenerative disease of musculoskeletal system. The total number of patients with OA disease has exceeded 250 million in the world, and the number of patients is still rising rapidly and continuously^1 2^. With the aging of our society, the incidence rate of osteoarthritis is increasing, which will become the most important factor leading to the dysfunctions of the elderly and seriously affect the quality of life of the patients^3 4^. At present, the clinical therapeutic effect of OA is still not ideal. Therefore, it is urgent to strengthen the research on relevant aspects so as to provide new theoretical support for the prevention and treatment of OA.

OA is a total joint disease involving cartilage, subchondral bone and synovium^5^. Among them, cartilage injury is the core pathological change of OA and the basic element of OA disease diagnosis. A variety of biological factors such as aging, inflammation and mechanical stimulation will destroy the homeostasis of chondrocyte, and then aggravate OA progression^6-8^. The articular cartilage consists of chondrocytes and extracellular matrix, which mainly depends on the infiltration of joint fluid to maintain nutrition. As there is no distribution of blood vessels, nerves and lymphatic vessels, the repair and regeneration ability of articular cartilage after injury is very poor^9^. Until now, there are no effective measures to protect OA chondrocytes from degeneration. More studies on the homeostasis maintenance of chondrocytes in the process of OA are needed to provide a new potential target for this disease.

Extrachromosomal circular DNA (eccDNA) refers to the circular DNA derived from genomic DNA but free of chromosomes. Recent studies have shown that eccDNA is extensively expressed in different tissues and may be involved in varied physiological and pathological processes including immune, inflammation, aging and tumors^10-13^. However, the expression of eccDNA in cartilage tissue and its pathological significance in OA are still unknown. Here, we extracted eccDNA from total knee arthroplasty (TKA) donors and firstly investigated the characteristics of eccDNA in articular cartilage using cir-DNA sequencing, and discussed the potential roles of eccDNA in OA.

## Methods

### Human Cartilage Tissue Collection and Isolation of Chondrocytes

Human cartilage was obtained from OA patients (n=10, aged 69.3 ± 6.53 years, four male and six females). Subjects with tumors, diabetes or other severe diseases in the last 5 years were excluded. Human articular cartilage explants were cultured according to the previously described methods^14^. Briefly, freshly obtained germ-free human cartilage was washed with PBS and cut into 4-mm-diameter blocks by the bistoury. Subsequently, the bocks were digested in high-glucose DMEM supplemented with 0.2% type II collagenase for 6h. Then the digestion suspension was collected and centrifuged at 400g for 5min following the filtration with a 40 μm cell strainer. Finally, the sediment (primary chondrocyte) was resuspended with high glucose DMEM containing 10% FBS and cultured in a 5% C02 incubator.

### Circle-Seq

The method of Circle-Seq Chondrocyte eccDNA was optimized based on the description of Henrik Devitt Møller et.al, including of multiple steps showed below: 1) Total DNA was extracted using QIAamp DNA Mini Kit (Qiagen); 2) Then, total DNA were alkaline treated to separate chromosomal DNA, lipids, and protein by rapid DNA denaturing–renaturing, followed by column chromatography on an ion exchange membrane column (Plasmid Mini AX; A&A Biotechnology); 3) Remaining linear DNA was removed by exonuclease (Plasmid-Safe ATP-dependent DNase, Epicentre), assisted by rare-cutting endonuclease MssI (only support Homo sapiens) that digested mitochondrial circular DNA (mtDNA, 16 kb) and made additional accessible DNA ends for exonuclease; 4) EccDNA-enriched samples was used as template for phi29 polymerase reactions (REPLI-g Midi Kit) amplifying DNA; 5) Amplified circular DNA was cleaned (AMPure XP beads) and sheared by sonication (Bioruptor) to average fragment size of 200 –300 bp; 6) Libraries for next-generation sequencing were prepared using the NEBNext Ultra DNA Library Kit for Illumina according to the manufacturer’s protocol (New England Biolabs), and subjected to sequencing on Illumina Novaseq 6000 using PE150, collecting up to 80 million paired-end reads/sample.

### EccDNA Mapped and identified Method for Circle-Seq Data

The sequencing reads from Circle-Seq were aligned to reference genome sequences (GRChg38) using the BWA program (Burrows-Wheeler-Alignment Tool, https://github.com/lh3/bwa) to dig eccDNA originated from the chromosome. Subsequently, the candidate reads were sorted, labeled, and ranked by Circle-Map to detect and identify eccDNA refers to the practices of Henrik Devitt Møller.

## Results

### cir-DNA sequencing revealed the landscape of eccDNA in articular cartilage from OA patients

Firstly, we harvested articular cartilage (n=10) from OA patients post-TKA operation (supplemental Figure 1A), and collected eccDNA of chondrocytes for cir-DNA sequencing (**Figure 1A** and supplemental Figure 1B). The quality of cir-sequencing output reads was evaluated as excellent (supplemental Figure 1C and D). Then we used BWA software to perform genomic alignment of reads with GRch38. For mapped reads, we utilized Circle-Map software for eccDNA identification and detected 39,342 eccDNA (detected eccDNA) (**Figure 1A**). Next, we developed a filter standard of split reads> 1 to identify the detected eccDNA, and finally 15121 eccDNA (identified eccDNA) were obtained (**Figure 1 B**). Subsequently, we analyzed the size of the identified eccDNA in Figure 1 B. We found that the size distribution of eccDNA in chondrocytes was between 0.1 kb and 10 kb, which was similar to the size distribution of eccDNA in muscle tissue but was distinct from that of serum (<1kb) and of tumor cells(1kb-1mb) (**Figure 1 C**). As to the distribution of eccDNA on chromosomes, the results uncovered that chondrocyte eccDNA distributed in all other regions except for Y chromosome and the short arm of chromosome 13,14,15, which was similar to that in Hela cells, but a bit different from that of glioma cells and yeast cells (**Figure 1 D**). In addition, previous studies have demonstrated that eccDNA may derive from repetitive sequences on the genome^15^. Indeed, we found that most of the reads corresponding to eccDNA of chondrocytes matched the repetitive sequences of the genome, including long interspersed nuclear elements (LINEs, 19.2%), short interspersed nuclear elements (SINEs_Alu, 18.8%), long terminal repeat (LTR, 11.3%), and a small number of telomere sequences (**Figure 1 E**), which was partly the same with muscle tissue, but different from plasma^16 17^. We further analyzed the potential roles of eccDNA in cartilage. Excluding the overlapping parts, 8915 genes were annotated, which included mRNA (6468), lncRNA (1075)), multipletype (697), pseudogene (219), miRNA (1), and others (455) (**Figure 1F**). These data indicate that articular cartilage contains an abundance of eccDNA, whose size range and chromosome origin may be relatively specific.

**Figure 1.**
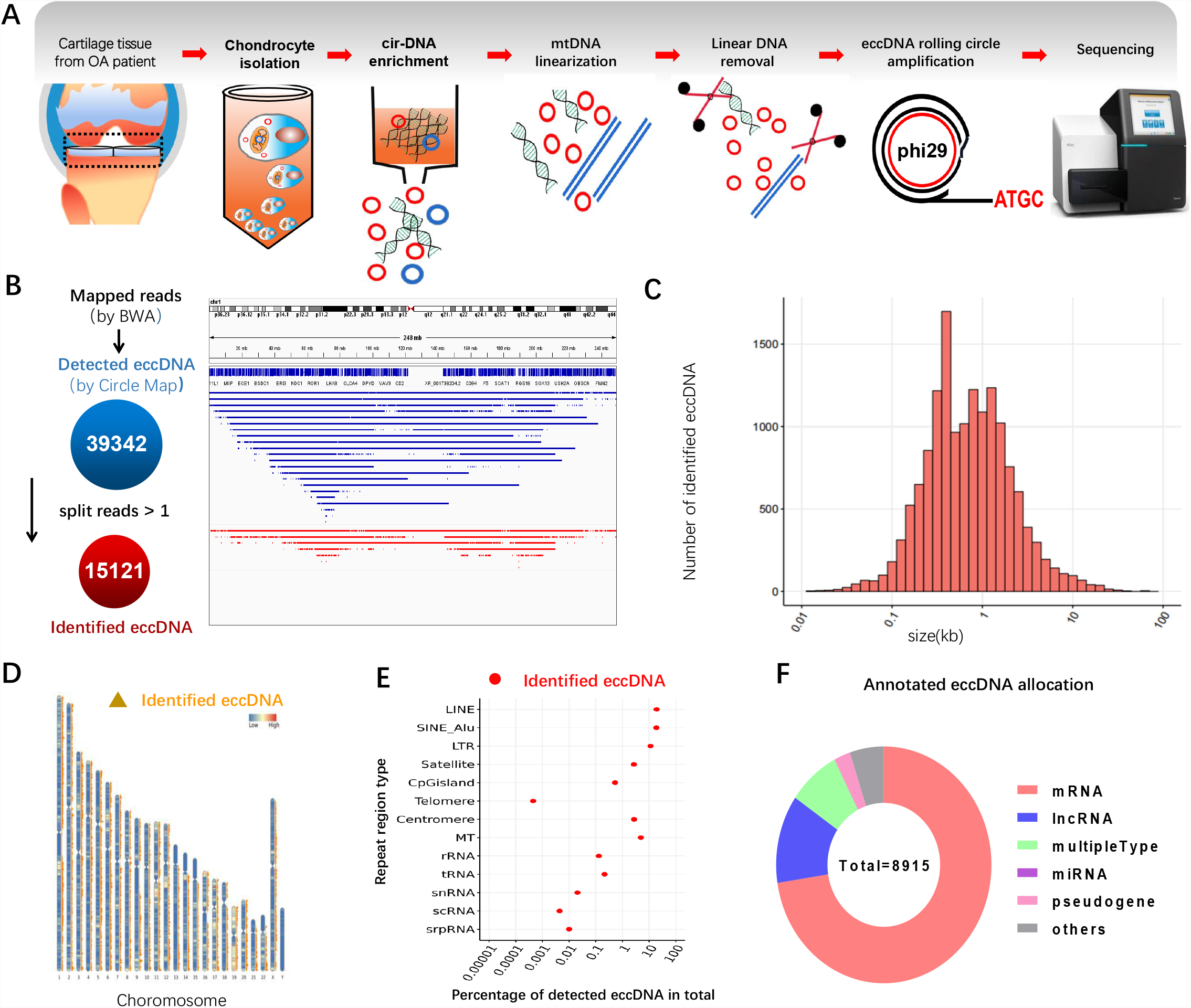

### OA-related transcription factors that involved in eccDNA cyclization

Previous studies have suggested that the junction point of eccDNA and its nearby area played an important role in eccDNA formation. Therefore, we separately extracted the 200bp sequence from upstream and downstream of the junction point start site or the stop site, combing with HOMER software to blast the extracted sequence with the known motif and predict the de novo motif (**Figure 2A**). To explore the role of the transcription factor in eccDNA formation and effects, we performed transcription factor-based enrichment analysis on mapped known motifs and de novo motifs respectively, and extracted the TOP20(based on p-value) collection of the two outputs (**Figure 2B**, supplemental table 1 and 2). Interestingly, most of the transcription factors with high p-value were highly related to the development and degeneration of cartilage, such as MYB, STAT3, NKX, NRF2, ZNF215, FOX. Moreover, some transcription factors including KLF4, NRF2, STAT3 and NF-kB have been proved to be the key pathogenic factors in OA (**Figure 2B**). These results indicate that OA-related transcription factors may be involved in the formation of eccDNA in chondrocytes.

**Figure 2.**
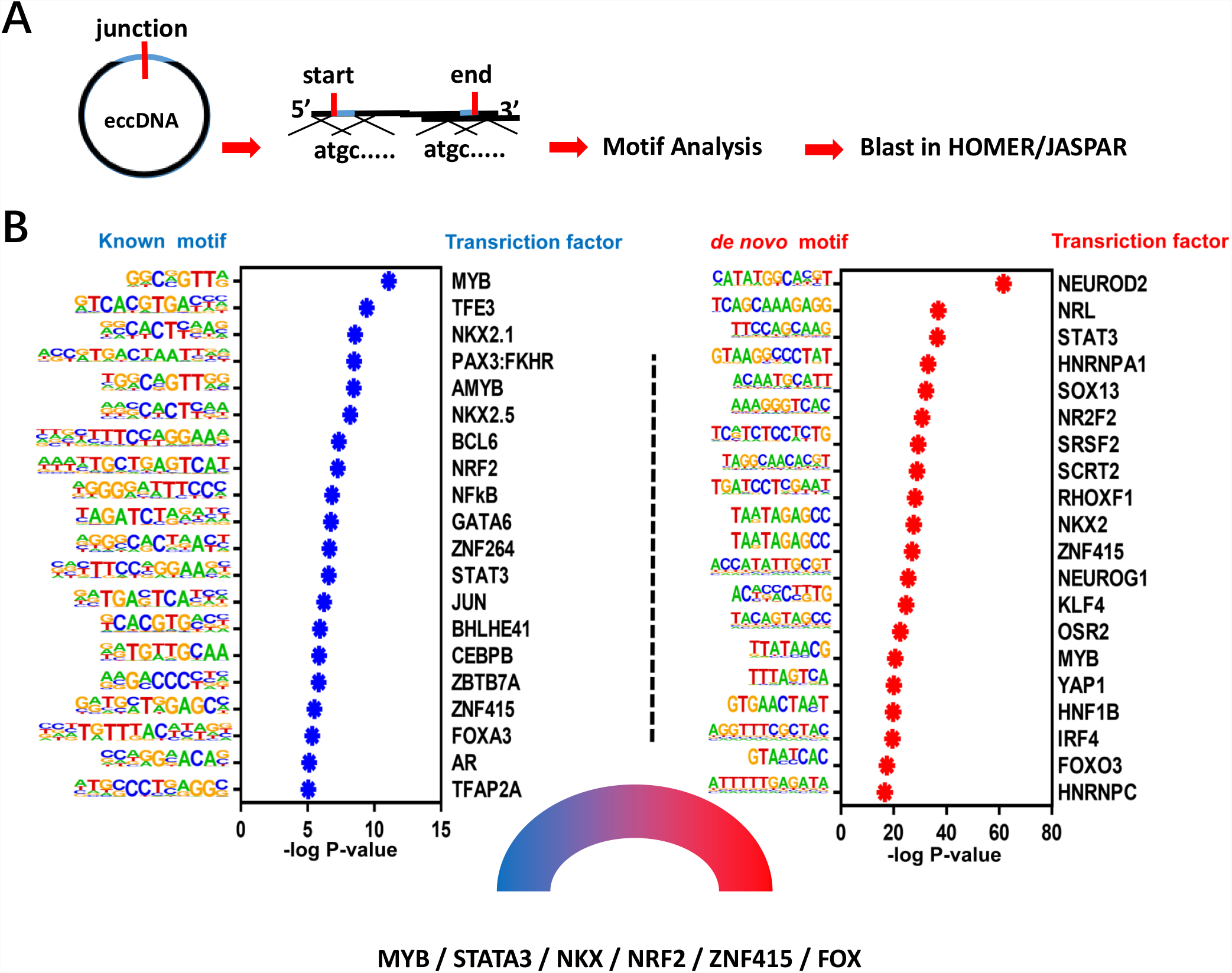

### The GO/KEGG pathways analysis for the eccDNA in OA

Next, we performed GO analysis and KEGG analysis of the above genes (**Figure 3A-B**). The data suggests that eccDNA may be involved in the pathophysiological regulation of degenerated chondrocytes in morphology, cell polarization, cell-ECM interaction, etc. From the top enriched terms, we found that abundant genes were involved in anabolism and catabolism, senescence, and inflammation of chondrocyte, such as ACAN, MMPs, ADAMTSs, IL1B, and so on (**Figure 3C**), indicating that eccDNA may participate in the regulation of cartilage homeostasis. Moreover, GO and KEGG analysis both revealed that the target genes of eccDNA were significantly enriched in nervous system-related pathways, such as axon guidance, synapse organization, suggesting that a potential role of eccDNA in joint pain of OA. Besides, we sorted the eccDNA according to circle score to analyze the genes annotated by TOP20 eccDNA and found that these genes are mainly related to the formation of nerve synapses (CNTN4, CABLES), cell stress response (SGK1), chromatin stability and DNA damage repair (NT5DC1, PDS5A, STK11), protein degradation and reparation (FBXW7, PCMT1, UBXN8), transcription inhibition (CBFA2T2, NCOR2), cell adhesion and skeletal remodeling (MAGI1, ARHGEF26) (**Figure 3C-D**). Furthermore, certain eccDNAs that potentially targeted OA-related genes from Figure 1J and 1K were analyzed and displayed (**Figure 3E**).

**Figure 3.**
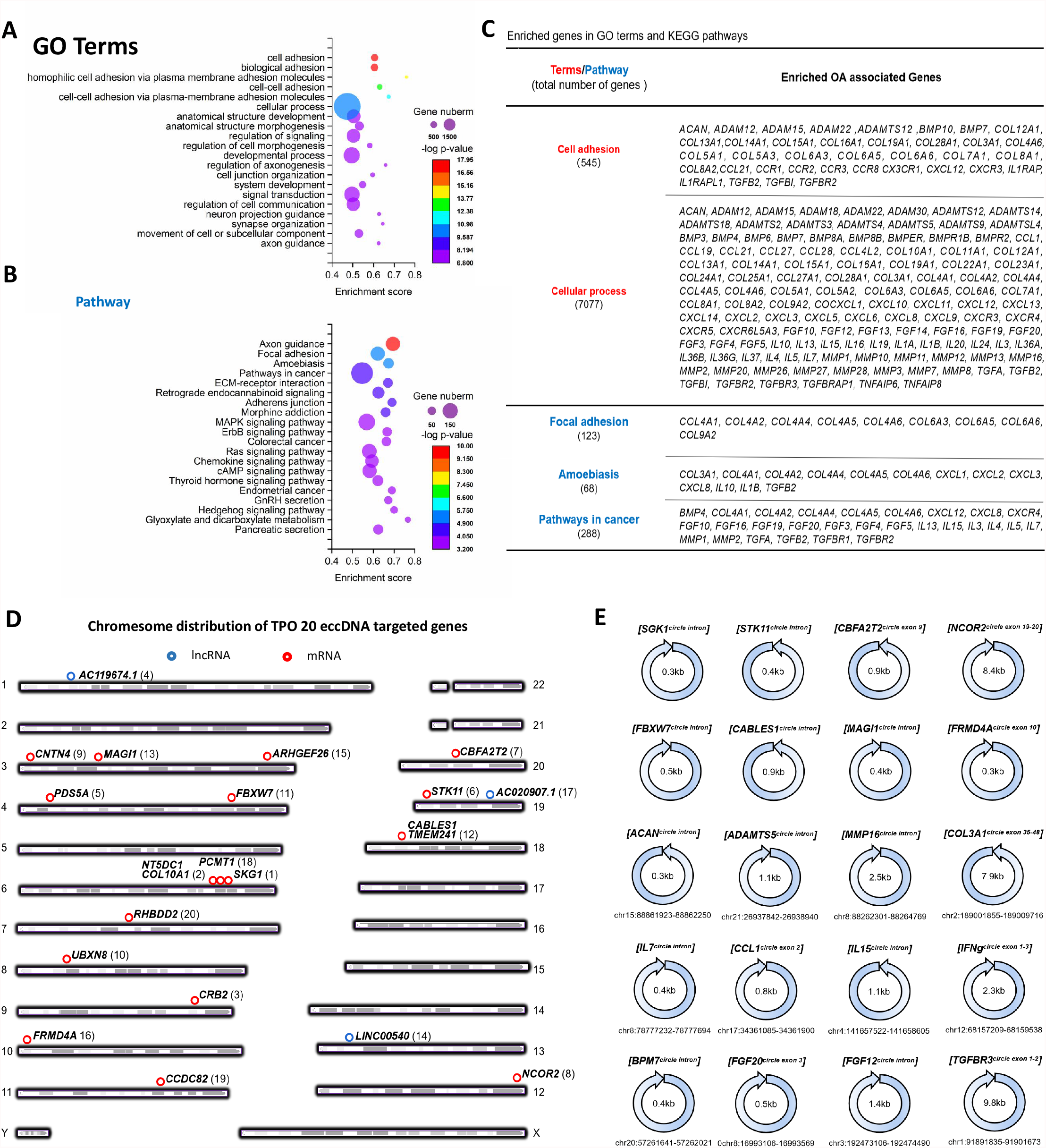

## Discussion

In recent years, more and more studies have shown that eccDNA is closely involved in the occurrence and development of several diseases such as cancer^13 18 19^. Turner, K. M. et al. reported that eccDNA was found in nearly half of human cancers and its frequency varied by tumor type^13 18 19^. They also revealed that driver oncogenes in eccDNA were amplified commonly, which contributed to the transcription of oncogenes out of chromosomes^20^. Wu, S. et al. investigated the structure of eccDNA by integrating ultrastructural imaging, long-range optical mapping and computational analysis of whole-genome sequencing, and their data suggested that oncogenes encoded on eccDNA increased copy number with high transcription levels^21^. In addition, eccDNA displays significantly enhanced chromatin accessibility than typical of chromosomes and has more ultra-long-range interactions with active chromatin ^21^, indicating it possesses powerful transcriptional capacity potentially. Koche, R. P. and their colleagues also found that cancer-causing lesions can increase the formation of functional eccDNA in neuroblastoma, which may play an important role in tumor progression^22^. Apart from tumor, eccDNA has also been related to aging process^23-25^. The accumulation of CUP1 eccDNA has unique-site specificity in the copper-induced aging model of yeast^24^. Furthermore, the heterogeneity of circular DNA that offers flexibility in adaptation is significantly diminished with age^25^. As aging is the main cause of OA, it is speculated that eccDNA may play an important role in the maintenance of chondrocyte homeostasis and the process of OA disease. Just like tumor cells, Gene transcription on eccDNA (not on Chromosomal DNA) may be actived in aged chondrocytes and be involved in OA process. As eccDNA comes from genome, it may regulate chondrocyte phenotypes by epigenetic means, such as affecting the replication and transcription of Chromosomal DNA. More studies are needed to investigate the roles and mechanisms of eccDNA on chondrocyte.

Synovium is an important part of the joint sliding system, which is composed of loose connective tissue. Its main functions include secreting synovial fluid, reducing joint surface friction, preventing joint adhesion, providing nutrients for articular cartilage, absorbing and swallowing various metabolites and cell fragments in the joint cavity, etc. When the joint is subjected to stress injury (such as degeneration, trauma, infection and rheumatism), the synovium will be stimulated to produce inflammatory reaction and participate in the repair of articular cartilage injury. It is found that chronic and low-grade inflammation is the main feature of OA synovitis, 6 which is closely involved in the formation of OA cartilage damage repair microenvironment. Therefore, trying to regulate the state of synovitis to reshape the microenvironment of OA joint, and then promote the damage and repair of chondrocytes, is expected to become a feasible strategy for clinical OA treatment. However, the formation mechanism of OA synovitis has not been very clear until now. Previous studies have suggested that the fragments from dead chondrocytes and degraded cartilage matrix can stimulate synovial tissue to produce inflammatory response and mediate the formation of chronic synovitis in the process of OA^26^. In addition, our group reported that the exosome-like vesicles from OA chondrocytes with inflammatory stimulation could increase the mature IL-1β production of synovial macrophages and aggravated synovitis in osteoarthritis^14^. Recently, Wang, Y. et al. revealed that eccDNAs are apoptotic products with high innate immunostimulatory activity^10^. Their data indicated that apoptosis inducers can increase eccDNA generation and eccDNAs can function as potent innate immunostimulants in a manner that is dependent on eccDNA circularity and the cytosolic DNA sensor Sting^10^. Therefore, we deduced that eccDNA from the apoptotic or senescent chondrocytes may participate in the formation of OA chronic synovitis via stimulating synovial inflammatory cells.

This study firstly mapped out the landscape of eccDNA in articular cartilage and preliminarily indicated the potential roles of eccDNA in OA, which may open up a new vision on the roles and mechanisms of eccDNA in joint diseases. Based on the biological characteristics of eccDNA^13 16 27-29^, we speculate that it may play a potential role in cartilage-related diseases, such as OA. The roles and mechanisms of eccDNA in joint diseases need further research. In addition, the formation mechanisms of eccDNA in chondrocytes and its changes in the pathological process are also worthy of further attention.

## Supporting information

supplemental table 1-Homer Known Motif Enrichment Results

supplemental table 2-Homer de novo Motif entichment Results

## Contributors

ZN, LK and YC conceived and designed the experiments. WX, TZ, YG, YW, JY, JY and LS performed experiments. YX, JL and XL provided expert advice. All authors analysed the data. WX, LK, XL and ZN wrote the paper.

## Ethics approval

All procedures and protocols were approved by the Ethics Committee with informed consent obtained (2018128) and carried out in accordance with standard operative procedures.

