## Supplementary figures and images for "cir-DNA sequencing revealed the landscape of extrachromosomal circular DNA in articular cartilage and the potential roles in osteoarthritis"

### Supplemental Figure 1

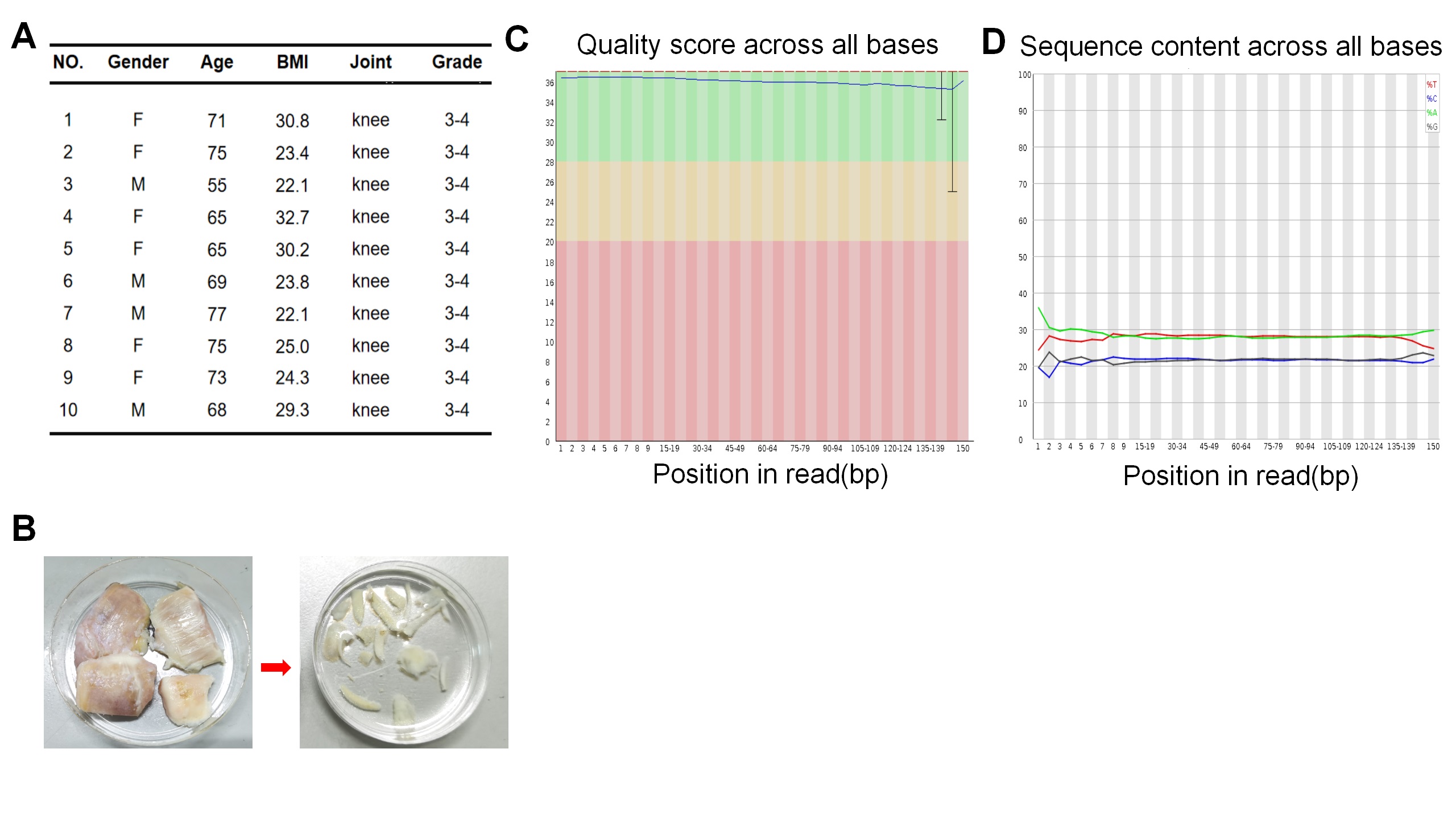
