## supplemental Figure legend for "cir-DNA sequencing revealed the landscape of extrachromosomal circular DNA in articular cartilage and the potential roles in osteoarthritis"

**Figure 1** The landscape of extrachromosomal circular DNA in articular cartilage. (A) Overview scheme of chondrocyte cir-DNA sequencing. Chondrocytes were isolated from Cartilage of OA patients (n=10) performed TCK and were put in a pool for sequencing. (B) eccDNA detected strategy. Output sequencing reads were mapped to GRCh38 by BWA, mappable reads were detected by Circle Map (blue) and filtrated based on split read >1(red). (C) Chondrocyte eccDNA distribution based on size. (D) Genomic coverage of chondrocyte eccDNA. (the bar is for chromosome, the blue and the red are for the density of genes; the orange peak is for eccDNA, peak height is for the density of eccDNA ). (E) The distribution of eccDNA on repeat regions from a genome. (F) The allocation of eccDNA based on target genes types(eccDNA were annotated with our scripts and bedTools, multiple types are for eccDNA were annotated more than 2 gene types).

**Figure 2** The potential transcription factors that participate in the formation of eccDNA. (A) The mode diagram of functional analysis for eccDNA cyclization from eccDNA junction breakpoint adjacent region (expand to 200bp upstream and downstream). (8) The Top 20 motif corresponding transcription factors and the OA-related ones were listed.

**Figure 3** The pathway and target gene analysis for eccDNA suggested the potential functions in OA. (A) TOP 20 gene ontology(GO) terms in eccDNA targeted GO analysis. (B) TOP 20 pathways in eccDNA targeted genes Kyoto Encyclopedia of Genes and Genomes(KEGG) analysis. (C) The enriched OA associated genes that eccDNA target in GO terms and KEGG pathways. (D) The chromosome distribution of TOP 20 eccDNA target genes according to circle score(circle score was according to mapping quality, fragment length, and split reads; number is for the ranking of genes). (E) Diagram of representative eccDNA for OA pathology.

**Supplemental Figure 1** (A) Information of OA patients. (B)Original cartilage samples from OA patients and diced cartilage blocks for chondrocyte isolation. (C) A quality score of reads output from cir-DNA sequencing. (D)Bases content of reads output from cir-DNA sequencing.
